# Adapting to changing methodology in a long-term experiment

**DOI:** 10.1101/2023.12.07.570659

**Authors:** Katherine McNamara Manning, Julia Perrone, Stephanie Petrycki, Douglas A. Landis, Christie A. Bahlai

## Abstract

Long-term experiments are critical for understanding ecological processes, but their management comes with unique challenges. As time passes, projects may encounter unavoidable changes due to external factors, like availability of materials, affecting aspects of their research methodology. At the Kellogg Biological Station Long-Term Ecological Research Site, one of many National Science Foundation-funded long-term research stations, a three-decade project recently experienced a supply-chain-induced change in insect sampling methodology in their lady beetle observation study. Since 1989, lady beetles (Coleoptera: Coccinellidae) have been sampled weekly over the growing season using yellow sticky cards. In 2021, the original sticky traps were discontinued by the manufacturer and replaced with a similar, but not identical trap. We conducted a 3-year study while the new traps were phased in to examine how the trap change would impact the observed biodiversity patterns at the site. We examined community metrics and individual taxa captures to examine within year and between year differences in performance between the card types. Overall, we noted several small but statistically detectable differences in capture patterns between the two trap types. After accounting for other sources of variation, we observed a difference in Shannon diversity of insects captured on the two card types, but not richness or abundance, in the overall insect community. Yet, these differences were dwarfed by the magnitude of difference observed between years within card types. For individual taxa, similar patterns held: between trap differences could be detected statistically, but the number of differences between capture rates of traps was less than the number of differences observed for the same trap, between years. Thus, we conclude that while subtle changes in methodology could impact data produced in long-term experiments, in this case the magnitude of this change is smaller than other factors such as time and plant treatment. However, if sustained changes in the capture rates of focal taxa are observed, future data users may use our observations to specifically quantify and correct for these shifting patterns related to the protocol change.

## Introduction

Long-term research is essential to understanding the ecology of systems (Cusser et al. 2021; Hughes et al. 2017; Lindenmayer and Likens 2009; Welti et al. 2021). However, with long-term research comes complex sampling histories and challenges that may alter certain aspects, such as changes of personnel or observers, changing goals or abilities of a project or site, unsuccessful planning, and even changes to sampling protocols due to availability of materials (Lindenmayer and Likens 2009; Vallecillo et al. 2020; Welti et al. 2021). In some cases, sampling protocol may be intentionally altered to respond to changing conditions at a site (e.g., to ensure adequate detection of taxa) (Blocksom, Emery, and Thomas 2009). Furthermore, long-term experiments are difficult to maintain in a traditional research environment. Most experiments are typically completed within a graduate degree or grant duration, leading to piecemeal management when funding must be compiled from multiple sources (Cusser et al. 2021). However, the National Science Foundation Long-Term Ecological Research Network (LTER) conducts long-term, place-based ecological research as a core infrastructure to support the ecological science community, providing the data collected at these sites to the public. This network, developed in response to arguments in support of long-term research (Callahan 1984), has since expanded to 28 terrestrial and aquatic sites globally, though most are in North America (Church et al. 2022). Because of the legacy of data and documentation made available in these studies, they provide an unprecedented opportunity to study methodological sensitivity in complex processes, like those examined in biodiversity studies. While experiments like these ideally buffer against management-imposed abrupt changes in protocol, some exceptions occur.

When methodology of a long-term experiment is changed, it is likely that these changes will cause some discontinuities in the data being recorded. Because each data point ultimately depends on the context in which it is being measured, any change to the measurement context between data points can change the signal being measured, resulting in the observation of different patterns even when data is taken at the same sites (Busse et al. 2022). Context-dependence is especially true of measurements taken to record biodiversity processes: because each datum is a culmination of multiple processes serving as a proxy for the desired measurement (i.e., ‘how many of this organism do we record with our method?’ vs. ‘what is the population density of this organism?’), some changes to biodiversity monitoring protocols will not be robust to these changes. For example, insect surveys can be extremely sensitive to the sampling methods used. Multiple studies have shown that sampling bees and other pollinating insects using different methods (for example: pan traps versus netting) captures different communities of pollinators (Berglund and Milberg 2019; Campbell et al. 2023; Grundel et al. 2011; Joshi et al. 2015; O’Connor et al. 2019; Prendergast et al. 2020; Wood, Holland, and Goulson 2015). Gardiner et al. (2012) examined the differences observed in lady beetle data collected by citizen scientists versus trained scientists, focusing on who collected the data rather than physical trapping methodology, but showing differences in richness and diversity estimates. McNamara Manning et al. (2022) compared trapping methodology for ground-dwelling arthropods and found that multiple biodiversity metrics differed between trap types. The sensitivity of biodiversity monitoring and observation methodology creates a challenge for both researchers maintaining long-term experiments and future researchers to synthesize results (Didham et al. 2020).

Although it would be ideal for long-term experiments to be maintained using identical protocols for their duration, changes may be required by circumstance, or may occur spontaneously (Lindenmayer and Likens 2009). How we manage a change during a long-term experiment might depend on whether we know the change is imminent or undetected until after the fact. Regardless of the source of the protocol change, when data produced by these experiments are a public resource, the transparency of those who maintain long-term experiments is critical. For instance, when six new sites were added to the grasshopper study at Konza Prairie, an LTER site in Kansas, USA, researchers noted that the intensity of sampling for the project was subsequently increased (Welti et al. 2021). When the change in sampling intensity was accounted for in models, they observed an annual decrease in grasshopper populations (Welti et al. 2020), but when sampling intensity was not accounted for in a subsequent synthesis, a false increase in the population was reported (Crossley et al. 2020). Also, at the Konza Prairie LTER, an unforeseen change in experimental structure occurred when samples from 1992-1995 were lost due to equipment failure before they could be identified, but they reported this loss publicly (Jonas and Joern 2007; Welti et al. 2021). Similarly, vegetation and vertebrates in Australian mountain ash forests were being monitored for over 25 years when wildfires burned about 50% of the long-term monitoring sites in the program. After consultation with collaborators, an adaptive monitoring approach was implemented to maintain the previous protocol while adding new study treatments to address newly generated research questions (Lindenmayer et al. 2011).

The Kellogg Biological Station (KBS) in southwestern Michigan, USA is an agricultural LTER site founded in 1987. One of the first experiments to begin at KBS was an insect monitoring experiment within the context of a large mixed-use cropping system. Lady beetles (Coleoptera: Coccinellidae) are important predators serving as biocontrol agents introduced into agricultural landscapes (Dixon and Dixon 2000; Gardiner et al. 2009). Starting in 1989 researchers began surveying lady beetle species with yellow sticky cards in the Main Cropping System Experiment (MCSE), made up of a variety of common crop replicates (Bahlai et al. 2013). The selection of yellow as the most attractive trap color was confirmed by a concurrent experiment (Maredia et al. 1992) The experiments goal was to monitor the lady beetle assemblages in the mosaic of crops (Colunga-Garcia, Gage, and Landis 1997; Colunga-Garcia and Gage 1998) and over time. Using legacy data from the first 30+ years of this experiment, insect community changes have been observed in multiple studies, with variation attributed to species interactions, density dependent factors, and environmental variability (Arnold et al. 2023; Bahlai et al. 2015; Hermann et al. 2016).

In 2021, the yellow sticky cards used since the inception of the experiment were discontinued by the manufacturer. While a very similar sticky card was identified to replace the former standard, they were not identical to the previous color. As insects are notorious for their sensitivity to color of traps (Holthouse, Spears, and Alston 2021; Muppudathi et al. 2018; Toler, Evans, and Tepedino 2005) and lady beetles in particular (Maredia et al. 1992), we conducted a study while the new cards were ’phased in’ to experimentally examine if, and how, these changes would impact the observations made at the site. We specifically ask: (1) Do the insect communities and individual taxa capture rates differ on different card types?; (2) If so, what community metrics differ and how do capture rates of individual taxa differ?; and finally (3) How do patterns of variation between trap types compare to other sources of variation affecting capture rates on traps, such as year? Because of the scale of this study (>250 samples per week during the growing season and an ongoing commitment to maintaining observations through the transition), the KBS lady beetle monitoring experiment provides an ideal environment to examine the sensitivity of insect communities and provide guidance for future scientists examining insect biodiversity data subject to sampling discontinuities or changes.

## Methods

This study documents the community of insects captured on yellow sticky cards at the Kellogg Biological Station Main Cropping System Experiment over a 3-year period, 2020-2022, as a new trap type was phased in. In this study the old, legacy yellow sticky cards and the new, replacement yellow sticky cards were compared using standard protocol within the same sampling season in 2021. To track typical between-year variation we used data collected from old sticky cards that were deployed during the 2020 season compared with that card type in 2021. In 2022, KBS implemented full sampling using the new sticky cards.

### Site description

Data in this study was collected at Michigan State University’s Kellogg Biological Station Long Term Ecological Research Station (KBS LTER) in southwestern Michigan, USA (42°24’N, 85°24’W). This study focused on an insect survey that began in 1989 as part of the MCSE and has included forest sites in the design since 1993 (Bahlai et al. 2013; 2015; Landis 2020). While lady beetle populations were the main focus of this insect survey, several other non-lady beetle predators were also recorded at various times in the duration of the survey (Colunga-Garcia, Gage, and Landis 1997; Colunga-Garcia and Gage 1998; Hermann et al. 2016). See https://lter.kbs.msu.edu/research/long-term-experiments/main-cropping-system-experiment/ (MCSE) and https://lter.kbs.msu.edu/%20research/long-term-experiments/successional-and-forest-sites/ (forest sites) for a detailed experimental design.

The KBS LTER MCSE (Main Cropping System Experiment) consists of seven treatments: an annual field crop rotation (maize, soybeans and wheat) under four levels of management intensity (conventional, no-till, reduced input, and biologically-based), alfalfa/switchgrass, poplar tree plantation, and early successional vegetation maintained by annual burnings. Forest sites consist of three treatments: conifer forest plantations, late successional deciduous forest fragments, and successional forests arising spontaneously on abandoned agricultural land. Each treatment was replicated six times for MSCE treatments and three times for forest treatments, each replicate had individual plot sizes of 1 ha. Observations were taken from five permanent sampling stations within each treatment-replicate combination (Arnold et al. 2023; Bahlai et al. 2013; 2015; Maredia et al. 1992). This experiment continues to date.

### Field and laboratory methods

Since 1989, un-baited, two-sided, yellow cardboard sticky cards (Pherocon, Zoecon, Palo Alto, CA; #FFE900) have been suspended at 1.2 m above ground-level at each sampling station (Bahlai et al. 2013) (**Figure 1**). In 2021, sticky cards from a new manufacturer, also un-baited, two-sided, yellow cardboard sticky cards (Pherocon, Trécé, Adair, OK; #FFE000) were phased in, due to a discontinuation of the previously used traps (**Figure 1**). During this field season, researchers deployed these new cards at two of the five stations in each treatment-replicate combination (stations 2 and 4 within each plot). The other three stations used the old sticky card.

**Figure 1:**
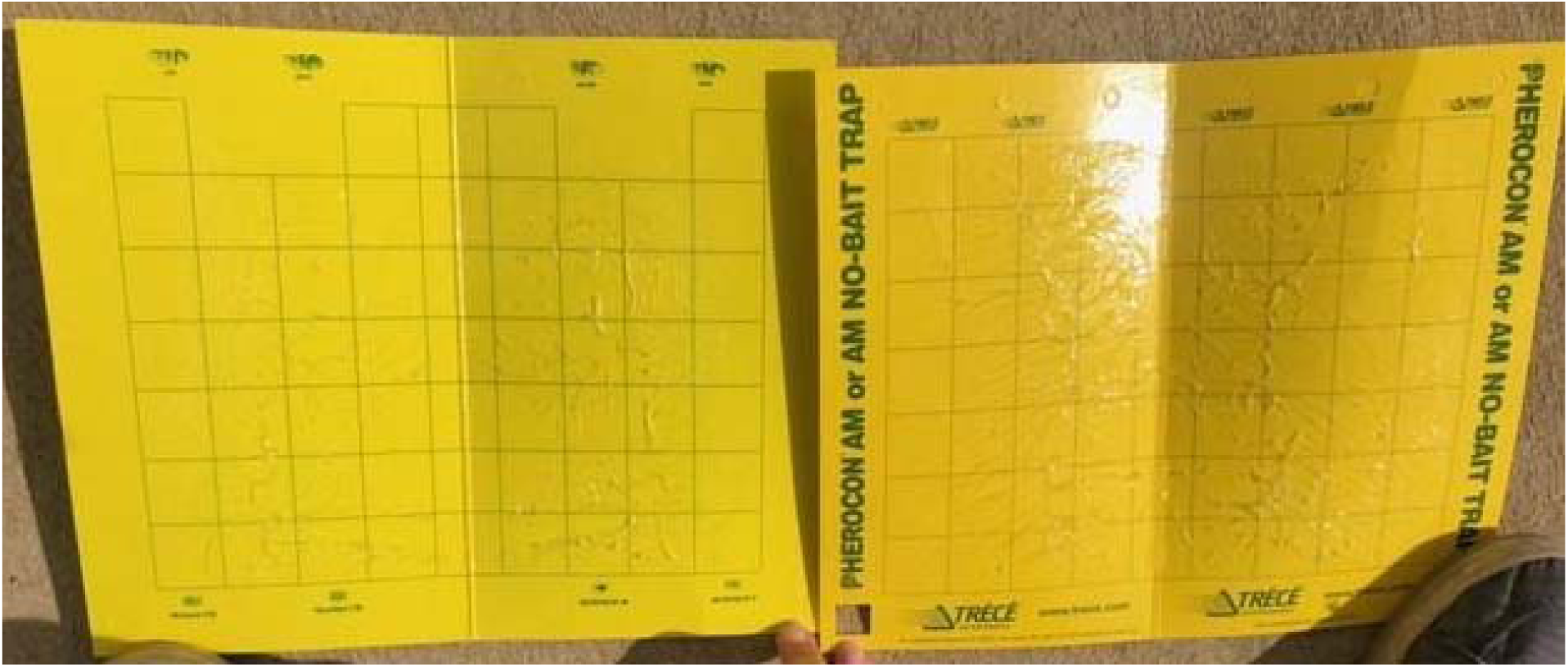
Yellow sticky cards. Left: Old card (Pherocon, Zoecon, Palo Alto, CA). Right: New card (Pherocon, Trécé, Adair, OK).

Each week, adult coccinellids and specified non-coccinellid predators on the sticky cards were identified using a pictorial key, counted, recorded, and then removed from sticky cards (Arnold et al. 2023; Bahlai et al. 2013). Sticky cards were replaced every two weeks from June to August of each year. In 2022, the same protocol was followed using only the new sticky cards (Pherocon, Trécé, Adair, OK).

The sticky cards collected in 2021 were shipped to Kent State University in Kent, OH, USA for further processing to increase the sensitivity of the comparison. Five additional taxa common to the site but not usually recorded by the monitoring program were identified on both the old and new sticky cards deployed: parasitoid wasps in the Chalcidoidea and Ichneumonoidea superfamilies, moths and butterflies (Lepidoptera), grasshoppers and crickets (Orthoptera), and hoverflies (Syrphidae).

The yellow sticky cards from the two manufacturers (Zoecon and Trécé) were compared using a spectrophotometer (Spectro 1, Variable, Chattanooga, TN). Scans of each sticky card produced hex numbers for each color, spectral curves, and comparisons of the colors under different light sources (incandescent (A), daylight - red shade (D50), daylight - neutral (D65), cool white fluorescent (F2), broad band white fluorescent (F7)).

### Statistical methods

All statistical analyses were completed using R 4.1.3 (R Core Team 2022). Insect counts were pooled by vegetation treatment replicate for counts of each taxa taken over a two week period. Taxonomic richness (number of taxa per sample), abundance (number of individuals per sample), and Shannon diversity index (Hill 1973) were calculated (Oksanen et al. 2022).

Generalized linear models (GLMs) were used to compare insect communities between old and new sticky cards within a year (2021), old cards between years (2020 and 2021), and new cards between years (2021 and 2022). The response variables examined were taxa richness, abundance, and Shannon diversity. Each model included a term for CARDxYEAR (type of card: old or new, and year deployed), WEEK (week of trap collection), TREAT (vegetation treatment), and TRAPS (number of traps per vegetation treatment replicate) as an offset, and followed the structure: Response variable ∼ CARDxYEAR + WEEK + TREAT + offset(log(TRAPS)). Poisson distribution was used for all models except Shannon diversity, which used Gaussian distribution.

GLMs were also performed for each individual taxa for each comparison: old and new cards within year (2021), old cards between years (2020 and 2021), and new cards between years (2021 and 2022). The models were similar to GLMs examining the entire insect community, except here each taxon collected was the response variable: Taxa ∼ CARDxYEAR + WEEK + TREAT+ offset(log(TRAPS)) and the data used contained only counts of that taxa and the environmental parameters. For all GLMs, the function ‘Anova’ from the car package (Fox and Weisberg 2019) used analysis of deviance to examine the effect of card type or year (CARDxYEAR) in each model.

To visualize the insect communities between comparisons we used non-metric multidimensional scaling (NMDS, with Bray-Curtis distance), computed using the vegan 2.5-7 package. An NMDS was run for each comparison: old and new cards within year (2021), old cards between years (2020 and 2021), and new cards between years (2021 and 2022).

Permutational multivariate analysis of variance (PERMANOVA) and analysis of multivariate homogeneity of group dispersions (BETADISPER) were performed following each NMDS (Oksanen et al. 2022).

Accumulation curves of insect taxa richness for old and new cards in 2021 were created using the BiodiversityR package (Kindt and Coe 2005). To estimate sampling efficiency for each card type, we used nonparametric Jackknife order 1 estimator to compare observed and estimated richness.

## Results

Over the three years we examined, 66,672 individual insects were collected and identified: 3,691 in 2020; 59,970 in 2021; and 3,011 in 2022. The five extra taxa we examined in 2021 added 54,517 individuals to that year’s total abundance, which was dominated by Chalcidoidea (parasitoid chalcid wasps). Two Coccinellid taxa that were found in previous years were not captured during these three years: *Adalia bipunctata* (L.) (Two-spotted lady beetle) and *Coccinella trifasciata* (L.) (Three-banded lady beetle). The most abundant lady beetle taxa during the study was *Coccinella septempunctata* (L.) (Seven-spotted lady beetle), followed closely by *Harmonia axyridis* (Pallas) (Multi-colored Asian lady beetle). Aside from Chalcidoidea, the most abundant non-coccinellid taxon was Lampyridae (fireflies) (**Table 1**).

**Table 1:**
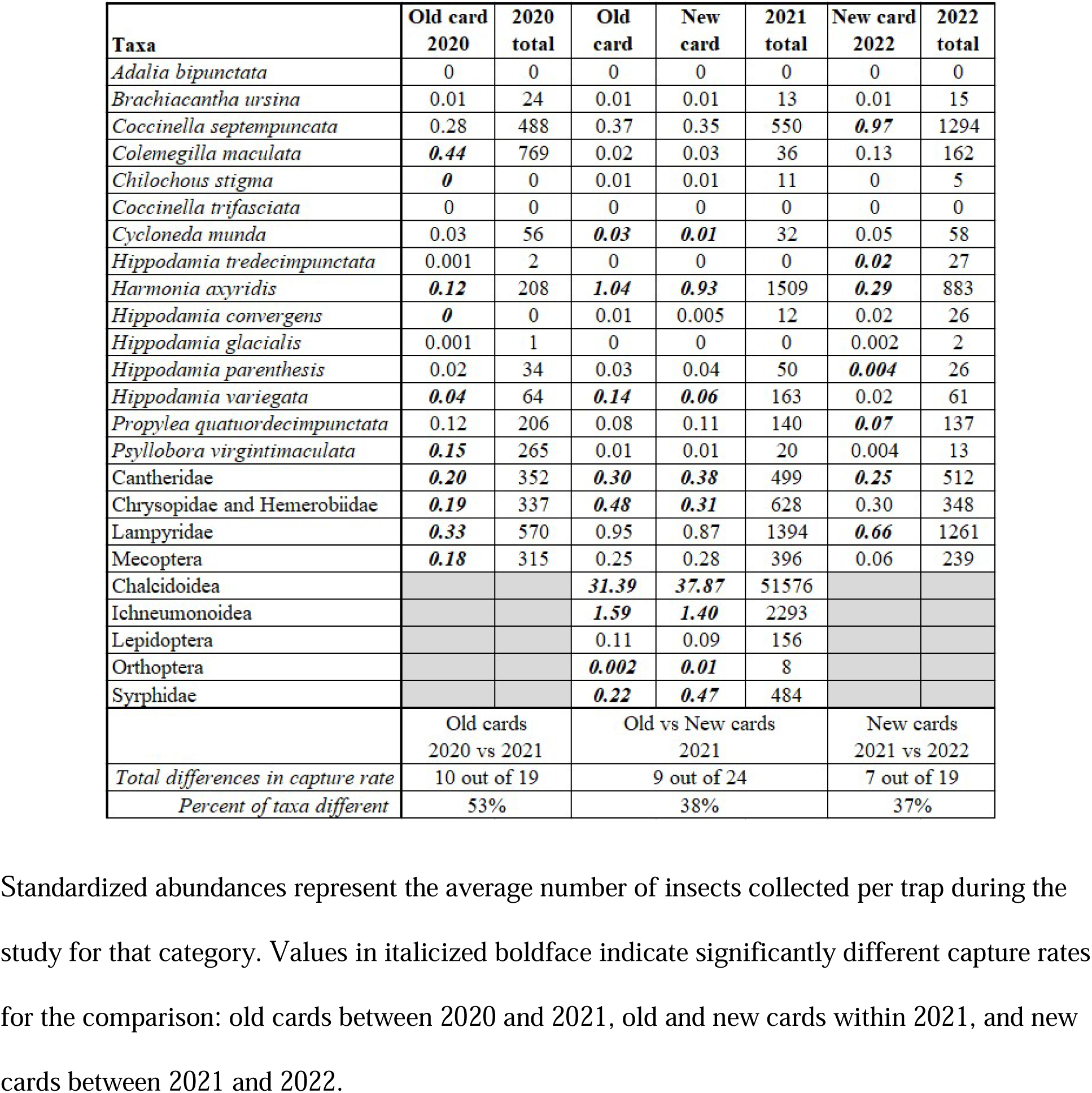
Insect abundances, total (raw) for each year and standardized for each card type per year.

The two sticky cards differed in several spectral properties measured. The color of the old yellow sticky card (Zoecon) was hex #FFE900 and had a reflectance peak at 530 nm, and again at 560 nm and 600 nm (**Figure S1**). The color of the new yellow sticky card (Trécé) was hex #FFE000 and had reflectance peaks at 530 nm, 560 nm, and 600 nm, but the peaks were greater at 560nm and 600 nm (**Figure S1**). There was an approximately 15% difference in reflectance between the card types at 490 nm and 530 nm, and 10% difference at 510 nm (**Figure S1**). When card types were compared under five types of light, three types of light showed perceptible differences between card types: daylight - red shade (ΔE = 2.90), daylight - neutral (ΔE = 3.14), and broad band white fluorescent (ΔE = 2.85) and two types of light did not show perceptible differences incandescent (ΔE = 2.24) and cool white fluorescent (ΔE = 1.84) (**Figure S2**).

### Within year variation

Comparing old and new cards deployed in 2021 using generalized linear models, we observed a significant difference in abundance (p<0.0001), but no difference in richness or Shannon diversity between card types within year (**Figure 2(b)**). Sampling week and plant treatment were significant in all models (**Table S1**). PERMANOVA testing for differences between card types following NMDS (stress = 0.23) indicated no difference in community composition (p = 0.30), and homogeneity of multivariate dispersion was assumed (**Figure 3(b)**).

**Figure 2:**
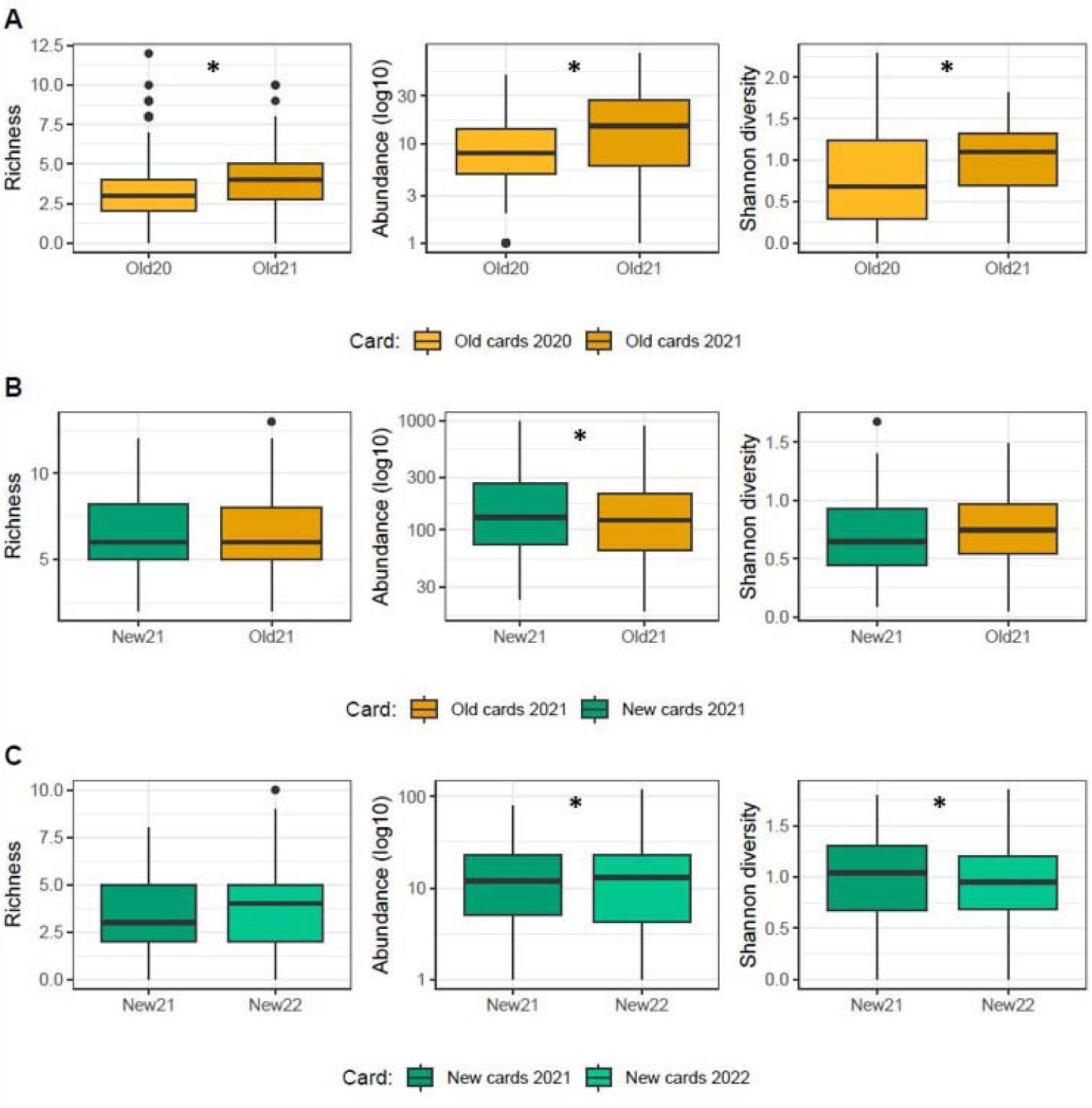
Richness, abundance (log 10), and Shannon diversity of insects for comparisons between (A) old cards in 2020 and 2021, (B) new and old cards in 2021, and (C) new cards in 2021 and 2022. Asterisk (*) represents significant difference (p < 0.05).

**Figure 3:**
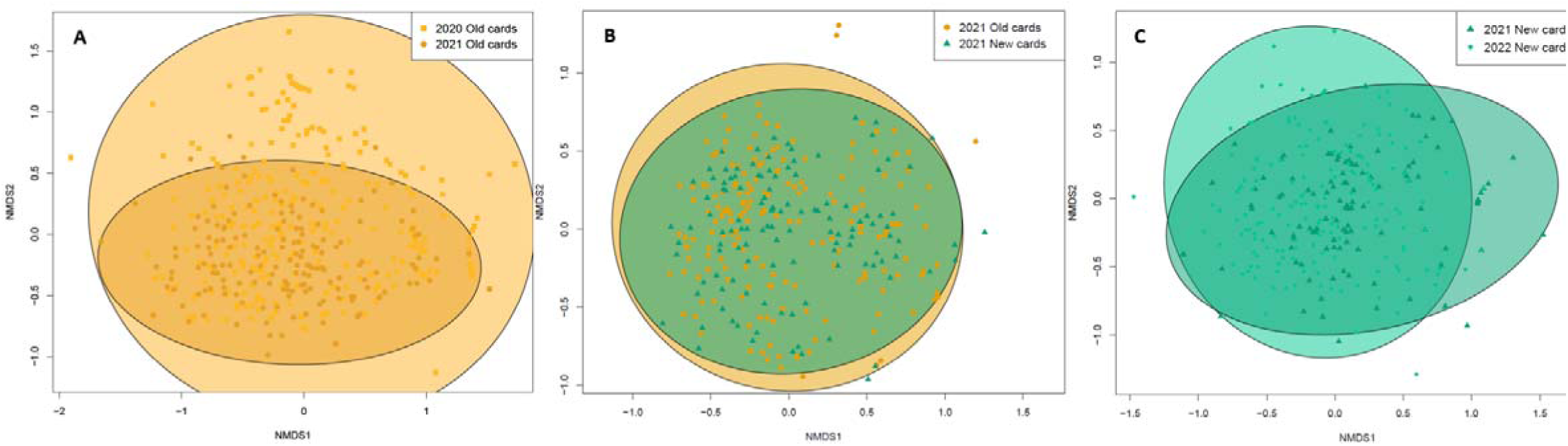
Non-metric multidimensional scaling displaying insect community composition for (A: stress = 0.18, p = 0.001) old cards in 2020 and 2021, (B: stress = 0.23, p = 0.30) new and old cards in 2021, and (C: stress = 0.23, p = 0.001) new cards in 2021 and 2022.

We surveyed for 24 insect taxa during 2021, and 21 of these taxa were collected on both the old and new sticky cards. When compared with first order jackknife richness estimates, capture efficiency of the old cards was 100% and of the new cards was 95%. There were fewer new cards deployed than old cards (i.e., they were deployed in a 2:3 ratio due to the five-subsample structure of the sampling design used in the MCSE). However, new cards followed an identical trajectory and thus it is likely that this difference is simply related to the number of samples taken (**Figure 4**).

**Figure 4:**
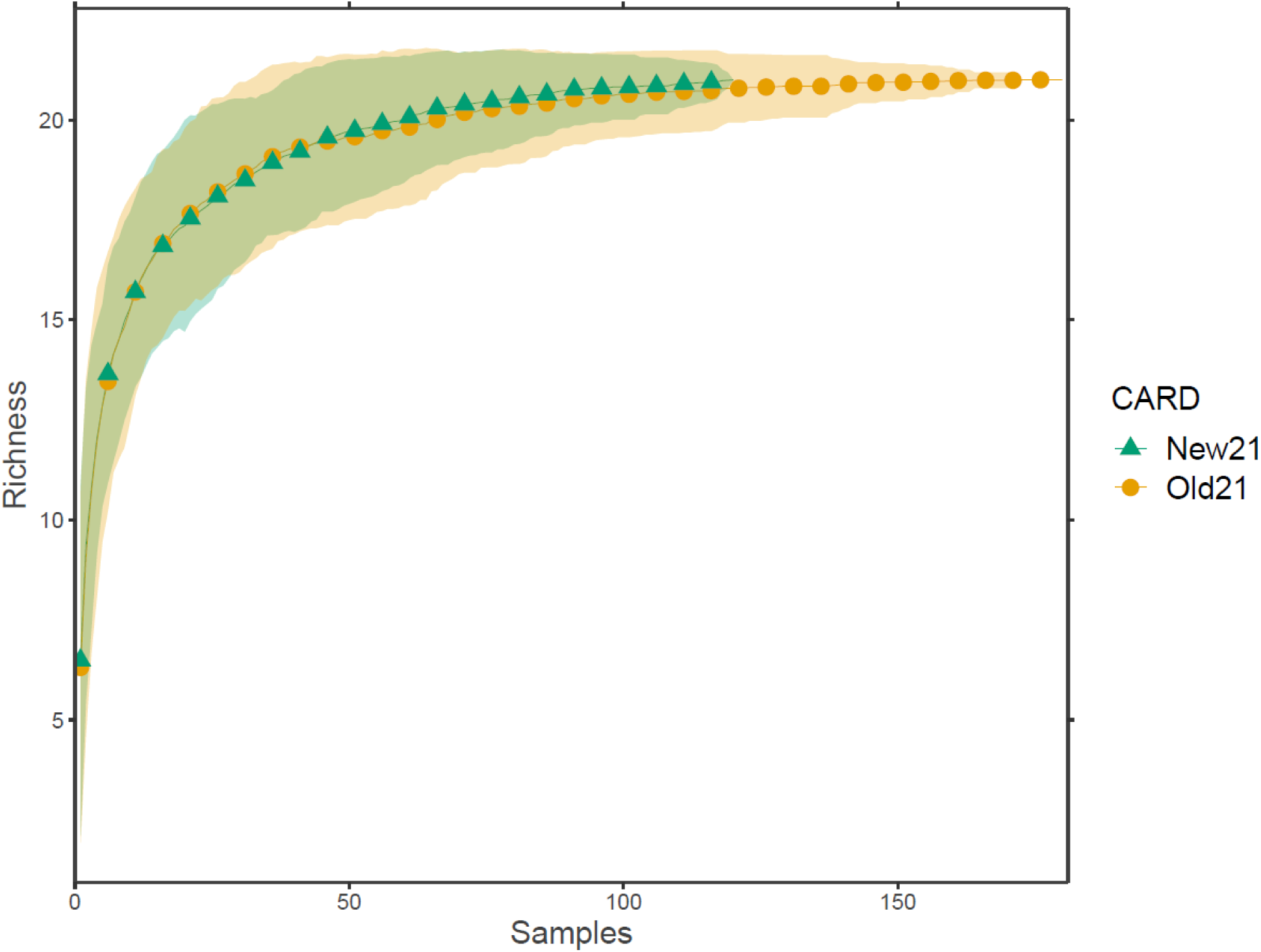
Accumulation curve for insect richness for old and new yellow sticky cards deployed in 2021. Shading represents standard deviation from the mean.

Within 2021, there were nine taxa where we observed statistical differences in captures per card between old and new cards. Five of these taxa were core taxa that were identified in the field per normal protocol: *Cycloneda munda* (Say), *Harmonia axyridis*, *Hippodamia variegata* (Goeze), Cantharidae, and Lacewings (Chrysopidae and Hemerobiidae), representing 26% of the 19 core taxa. The other four taxa with recorded differences in abundance in 2021 included four of the five additional taxa added in the laboratory with our additional processing: Chalcidoidea, Ichneumonoidea, Orthoptera, and Syrphidae. Altogether, these nine taxa were 38% of the 24 taxa surveyed for 2021 (**Table 1; Table S1; Figure S3**). Plant treatment was significant in all models, and sampling week was significant for all but two species (**Table S1**).

### Between year variation

Between old cards deployed in 2020 and 2021, we observed significant differences in richness, abundance, and Shannon diversity (p<0.0001) (**Figure 2(a)**). Sampling week and plant treatment were significant in all models (**Table S1**). PERMANOVA testing for differences in old cards between years following NMDS (stress = 0.18) indicated the communities captured differed significantly (p = 0.001) and homogeneity of multivariate dispersion could not be assumed (**Figure 3(a)**).

Between new cards deployed in 2021 and 2022, we observed significant differences in abundance and Shannon diversity (p<0.0001), but not for richness (**Figure 2(c)**). Sampling week was significant in all models, and plant treatment was significant for abundance and Shannon diversity (**Table S1**). PERMANOVA testing for differences in new cards between years following NMDS (stress = 0.23) indicated the communities captured differed significantly (p = 0.001) and homogeneity of multivariate dispersion was assumed (**Figure 3(c)**).

Comparing captures per card of individual taxa on old cards between 2020 and 2021, there were 10 taxa that differed in capture rate: *Coleomegilla maculata* (De Geer), *Chilocorus stigma* (Say), *Harmonia axyridis*, *Hippodamia convergens* Guerin*, Hippodamia variegata*, *Psyllobora virgintimaculata* (Say), Cantheridae, Lacewings: Chrysopidae and Hemerobiidae, Lampyridae, and Mecoptera. These 10 taxa represent 53% of the 19 taxa surveyed (**Table 1; Table S1; Figure S4**). Plant treatment was significant for all but three species, and sampling week was significant for all but four species (**Table S1**).

Comparing captures per card of individual taxa on new cards between 2021 and 2022, there were 7 taxa that differed in capture rate: *Coccinella septempuncata*, *Hippodamia tredecimpunctata* (L.), *Harmonia axyridis*, *Hippodamia parenthesis* (Say), *Propylea quatuordecimpunctata* (L.), Cantharidae, and Lampyridae. These 7 taxa were 37% of the 19 taxa surveyed (**Table 1; Table S1; Figure S5**). Plant treatment was significant for all but three species, and sampling week was significant for all but four species (**Table S1**).

## Discussion

Overall, comparisons between card type within a given sampling year had fewer differences than between year variation using cards of the same type, suggesting the effect that changing the trapping method has on the data is likely less than typical year to year insect community variations. The magnitude of the difference, when differences occurred, was usually much lower between trap types than between years. Plant community treatment and sampling week accounted for the bulk of the variation within individual taxa or community metrics. Given that the bulk of applications of these data focus on year-to-year trends or differences between plant community treatments, the change of cards will likely not lead to a fundamental discontinuity in taxa captured or biodiversity metrics observed within the data produced by this long-term experiment. However, for several taxa and metrics, statistically detectable differences did occur. Thus, our study provides context if major, systematic differences between captures and communities persist in subsequent years.

The within year comparison between the two card types suggested there was a difference in only one community metric: abundance. Mean abundance was higher on new cards (43.4 ± 17.6 on new cards and 36.9 ± 13.6 on old cards). However, card type only explains about 1% of the estimated deviance for abundance. On the other hand, almost all biodiversity metrics differed between years when trap type was held constant. When examining individual core taxa, that is, taxa identified for all three years of the study, we observed that fewer taxa had differences in capture rate between the two card types within year than the same card type between years, and the magnitude of the difference was generally smaller between traps than between years. Among the five core taxa with significant within year differences, capture rates were higher on the old cards for four of the taxa, suggesting the old cards may be generally slightly more attractive, or more efficient at capturing key taxa, assuming that actual populations of these taxa are relatively constant at this site.

In 2021, we chose five additional taxa to record on the sticky cards to provide additional statistical power to our comparison of the old and new card types. Aside from Orthoptera, which are herbivores (Joern 1979) commonly captured on these traps, we selected these taxa because they are important beneficial insects either providing biocontrol (Chalcidoidea, Ichneumonoidea, Syrphidae) or pollination services (Lepidoptera, Syrphidae) (Bonet 2009; Herrmann, Buchholz, and Theodorou 2023; Skevington et al. 2019). Given that these taxa are very commonly captured and known to be sensitive to environmental conditions (Benthall et al. 2022; Eckberg et al. 2015; Holthouse, Spears, and Alston 2021; Toennisson, Klein, and Burrack 2019), they provided an additional, more sensitive test beyond that which would be provided by simply examining the core taxa monitored at the site. Indeed, capture rates of four of the five additional taxa revealed a statistical difference between card types: the new cards appeared to be more attractive to Chalcid wasps, hoverflies, and Orthoptera, but less attractive to Ichneumon wasps.

Most studies of insect trap efficiency compare traps of different structures, colors or other major elements, and they almost uniformly find differences in trapping efficiency (Berglund and Milberg 2019; Boetzl et al. 2018; Busse et al. 2022; Campbell et al. 2023; Csaszar et al. 2018; Holthouse, Spears, and Alston 2021; Joshi et al. 2015; McNamara Manning, Perry, and Bahlai 2022; Muppudathi et al. 2018; Patrick and Hansen 2013; Prendergast et al. 2020; Shrestha et al. 2019; Toler, Evans, and Tepedino 2005; Work et al. 2002). This study is relatively unique in the subtlety of the comparison and was conducted in an existing experiment with high statistical power to test for small differences. We examined one type of trap, the yellow sticky card, but between two manufacturers, which varied in some of the physical aspects of the cards. Indeed, with spectrophotometer readings we confirmed that the two types of yellow sticky cards were different colors. Under neutral and warm daylight, in which sampling is conducted, the colors of the two cards had perceivable differences with the spectrophotometer and to the human eye. Although the cards had similar spectral reflectance curves, there were differences in percent reflectance, mostly from 490 – 530 nm. Color vision is used for finding food, shelter, mates, and other tasks, but varies among insects (van der Kooi et al. 2021). Though only one species of Coccinellidae was included, van der Kooi et al. (2021) found that their spectral sensitivity maxima were at 360 nm, 420 nm, and 520 nm, with 520 nm falling in the range we found the two sticky card types to differ in their percent reflectance (490-530 nm). Some of the other insects recorded also have spectral sensitivity maxima in that wavelength range: Ichneumonidae (530 nm), Orthoptera (Gryllidae ∼515 nm, Acrididae 430-515 nm), and Lampyridae 500-560 nm (van der Kooi et al. 2021).

Although we found relatively minimal differences between the two sticky cards when examined in the same sampling period, these differences occurred at a higher rate than what we would expect simply by chance alone, suggesting that, however subtle, differences do exist between the traps and their rates of insect capture. However, these differences are dwarfed by those associated with plant treatment and time (year and sampling week), making it unlikely to affect overall long-term trends in the data observed for the core taxa at this site. Indeed, other insect studies have noted more dramatic differences in comparisons of trap efficiency between methods. In another comparison experiment, trap color and type in bee monitoring was examined, finding the captures differed in abundance and richness, as well as that some bee subfamilies and tribes preferred different trap colors (Joshi et al. 2015). When the size of pitfall trap was altered, the community being captured changed, with larger diameter traps catching more arthropods, and more captured specimens belonged to common species (Work et al. 2002). Though meant to sample the same insect community, ground-dwelling arthropods, when pitfall and yellow ramp traps were compared, they captured significantly different richness, abundance, Shannon diversity, and evenness and had different community compositions (McNamara Manning, Perry, and Bahlai 2022).

## Conclusions

Changing the trap type led to small, but detectable changes in the insect community observed. Because insects are so sensitive to changes in monitoring, studies such as ours are vital to quantify impacts of methodological adjustments on on-going long-term experiments. In these cases, researchers who maintain long-term experiments face a profound responsibility to document and test changes in methodology or protocol, and most imperatively, be transparent about these changes to maintain the integrity of the project. We have demonstrated that even very subtle changes can lead to detectable changes, and thus we recommend, whenever possible, to incorporate a ‘phase in’ study so the magnitude of these changes can be estimated in the context of other variables that are more important to experimental goals. Thus, we conclude that even subtle changes in methodology could impact data produced in long-term experiments moving forward, and the key to ensuring continued integrity of the data is to document and quantify these changes. If managers see unexplained changes in the future of this experiment, they may look to this study for potential answers.

## Acknowledgements

We wish to acknowledge the research staff at the Kellogg Biological Station past and present, especially Stacey Vander Wulp, and the countless summer technicians who have maintained the experiment over the years. We thank Grace Chen for assistance with data entry, Morgan Hughes with assistance with insect processing, and Nick Haddad for ongoing support with long-term experiment maintenance. Support for this work was provided by the National Science Foundation Long-term Ecological Research Program (DEB 1832042) at the Kellogg Biological Station, NSF grants DBI 2045721 and DEB 2225092 to CAB, and Michigan State University AgBioResearch support to DAL. The long-term lady beetle experiment at KBS LTER was initiated in 1989 by Stuart Gage and Manuel Colunga-Garcia.

