## Supporting information for "Adapting to changing methodology in a long-term experiment"

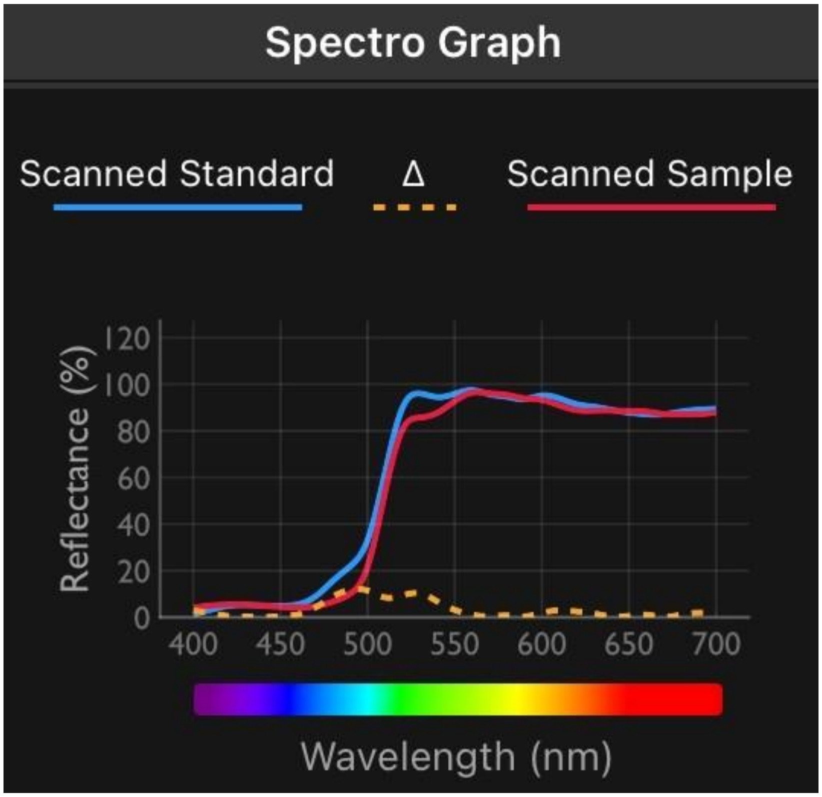


Figure S1: Spectral reflectance curves created with Spectro 1 spectrophotometer. Scanned standard (blue) represents the old sticky card (Zoecon) and scanned sample (red) represents the new sticky card (Trécé). Δ (dashed orange) represents the difference between the two types of sticky cards.

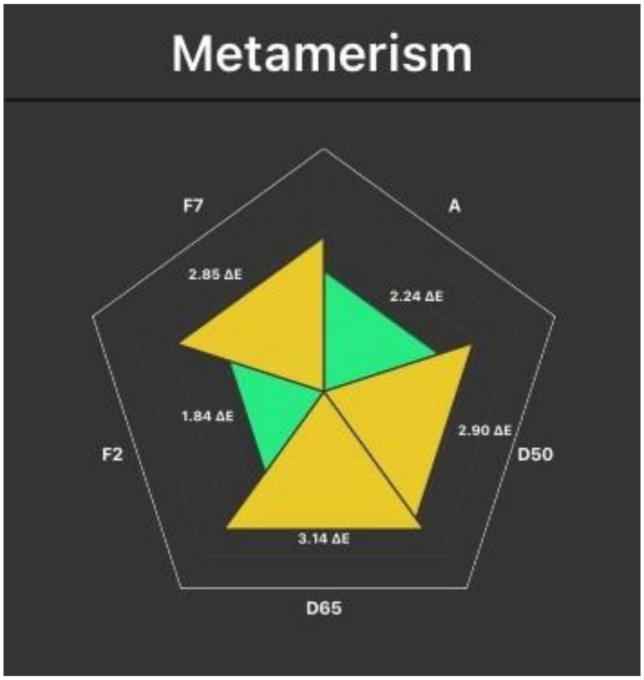


Figure S2: Metamerism alert figure created with Spectro 1 spectrophotometer. Card types were compared under five types of light: incandescent (A), daylight - red shade (D50), daylight - neutral (D65), cool white fluorescent (F2), broad band white fluorescent (F7). ΔE represents the difference between the old sticky card (Zoecon) and the new sticky card (Trécé) for each light type. Green triangles mean the difference between card types is not perceptible for that light type and orange triangles mean the difference between card types is perceptible for that light type.

Table S1: Estimated percent deviance explained by each variable (card type or year, sampling week, and plant treatment) for each model (richness, abundance, Shannon diversity for whole community, and capture rate of each individual taxa). Values in italicized boldface indicate significance of that variable within that model.

| | Within year | | | Old cards between years | | | New cards between years | | |
|---|---|---|---|---|---|---|---|---|---|
| | Card type | Sampling week | Plant treatment | Year | Sampling week | Plant treatment | Year | Sampling week | Plant treatment |
| Richness | < 0.1% | ***23%*** | ***14%*** | ***4%*** | ***19%*** | ***19%*** | 2% | ***35%*** | 4% |
| Abundance | ***1%*** | ***21%*** | ***40%*** | ***11%*** | ***38%*** | ***5%*** | ***2%*** | ***56%*** | ***6%*** |
| Shannon Diversity | < 0.1% | ***2%*** | ***95%*** | ***0.4%*** | ***9%*** | ***79%*** | ***11%*** | ***24%*** | ***41%*** |
| ABIPN | NA | NA | NA | NA | NA | NA | NA | NA | NA |
| BURSI | < 0.1% | ***18%*** | 15% | 1% | ***16%*** | ***32%*** | < 0.1% | ***24%*** | 13% |
| C7 | < 0.1% | ***41%*** | ***24%*** | 1% | ***38%*** | ***18%*** | ***8%*** | ***65%*** | ***14%*** |
| CMAC | 0.2% | ***8%*** | ***14%*** | ***17%*** | ***37%*** | ***29%*** | 11% | ***26%*** | ***14%*** |
| CSTIG | 1% | ***25%*** | ***43%*** | ***20%*** | 21% | ***34%*** | 19% | 21% | ***43%*** |
| CTRIF | NA | NA | NA | NA | NA | NA | NA | NA | NA |
| CYCSP | ***4%*** | 6% | ***1%*** | < 0.1% | ***17%*** | ***15%*** | 8% | ***22%*** | ***20%*** |
| H13 | NA | NA | NA | 7% | 31% | 35% | ***15%*** | ***28%*** | 7% |
| HAXY | ***0.1%*** | ***44%*** | ***30%*** | ***35%*** | ***19%*** | ***13%*** | ***18%*** | ***27%*** | ***23%*** |
| HCONV | 2% | ***19%*** | 16% | ***27%*** | 12% | 10% | 5% | ***46%*** | ***11%*** |
| HGLAC | NA | NA | NA | 7% | 31% | 35% | 9% | 28% | 30% |
| HPARN | 0.2% | ***10%*** | ***22%*** | 1% | ***8%*** | ***23%*** | ***1%*** | 12% | ***20%*** |
| HVAR | ***5%*** | ***25%*** | ***20%*** | ***13%*** | ***16%*** | ***15%*** | 8% | ***30%*** | ***22%*** |
| PQUA | 0.5% | ***18%*** | ***24%*** | 1% | ***21%*** | ***17%*** | ***2%*** | ***23%*** | ***17%*** |
| X20SPOT | 0.2% | 3% | ***27%*** | ***11%*** | ***16%*** | ***41%*** | 7% | 13% | ***33%*** |
| CANTHARID | ***0.5%*** | ***72%*** | ***15%*** | ***1%*** | ***51%*** | ***22%*** | ***1%*** | ***51%*** | ***14%*** |
| LAMPY | 0.1% | ***52%*** | ***8%*** | ***14%*** | ***36%*** | ***3%*** | ***1%*** | ***64%*** | ***3%*** |
| LCW | ***3%*** | ***21%*** | ***28%*** | ***8%*** | ***16%*** | ***31%*** | 7% | ***22%*** | ***15%*** |
| MECOP | 0.1% | ***22%*** | ***62%*** | ***0.5%*** | ***16%*** | ***62%*** | 10% | ***32%*** | ***44%*** |
| Syrphidae | ***6%*** | ***38%*** | ***17%*** | | | | | | |
| Ichneumonoidea | ***0.3%*** | ***11%*** | ***54%*** | | | | | | |
| Chalcidoidea | ***1%*** | ***21%*** | ***44%*** | | | | | | |
| Lepidoptera | 0.2% | ***18%*** | ***17%*** | | | | | | |
| Orthoptera | ***7%*** | ***29%*** | ***32%*** | | | | | | |

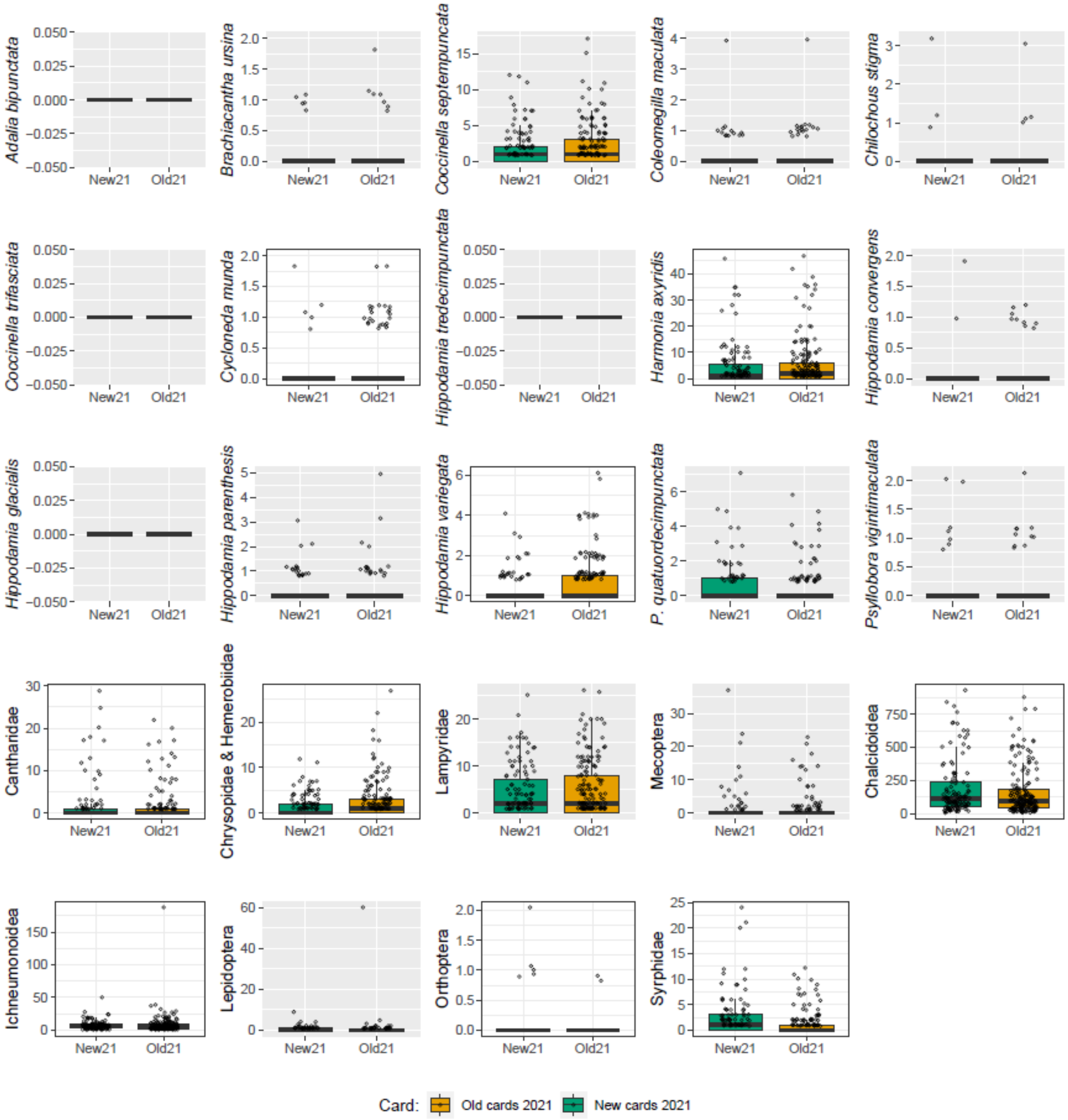


Figure S3: Captures of individual taxa for old and new cards in 2021. Plots with a white background indicate significant differences between card type ($p < 0.05$). Plots with grey backgrounds indicate no statistical difference in capture rates were detected in this taxon.

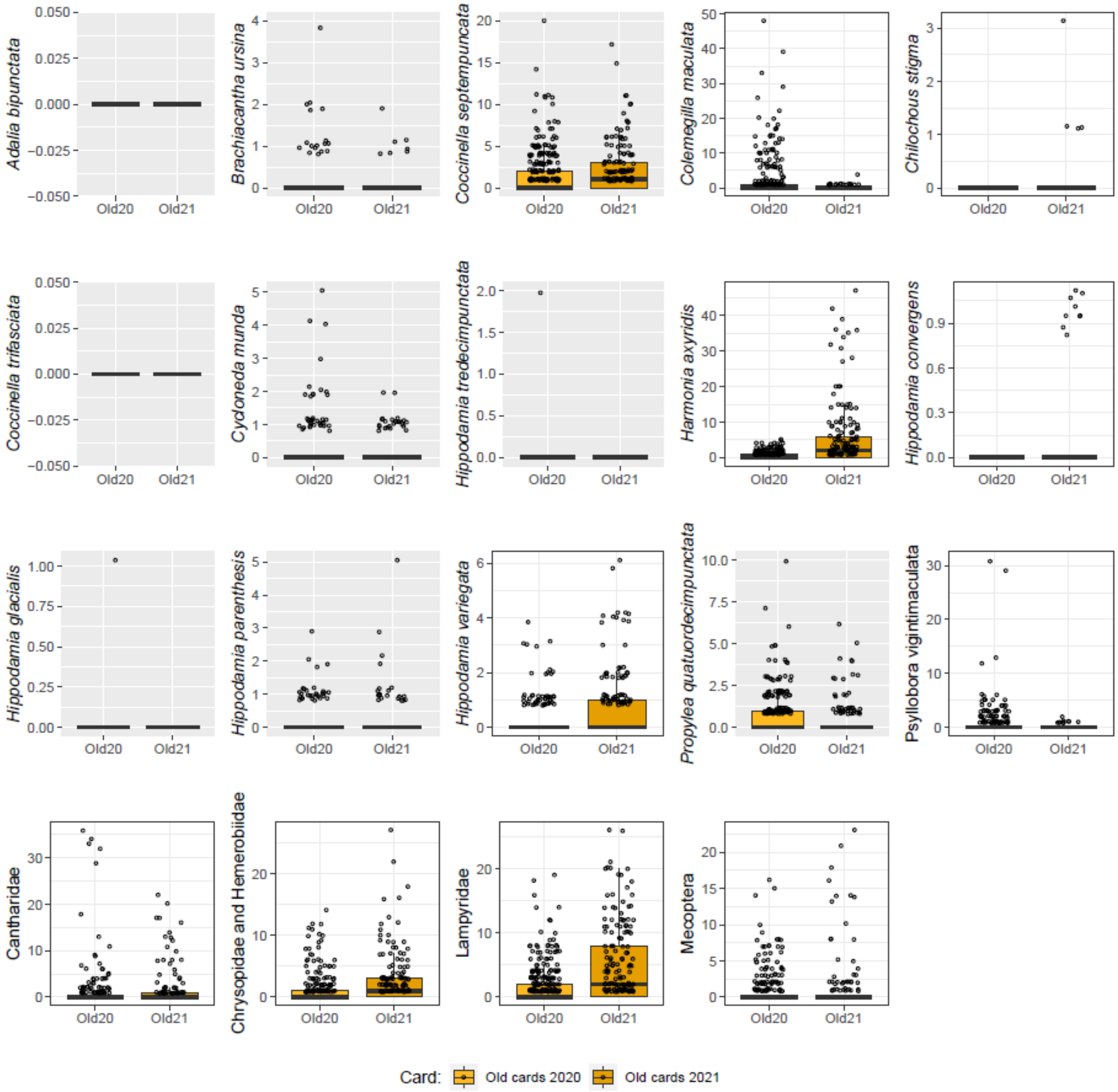


Figure S4: Captures of individual taxa for old cards in 2020 and 2021. Plots with a white background indicate significant difference in capture rates between year ($p < 0.05$). Plots with grey backgrounds indicate no statistical difference in capture rates were detected in this taxon.

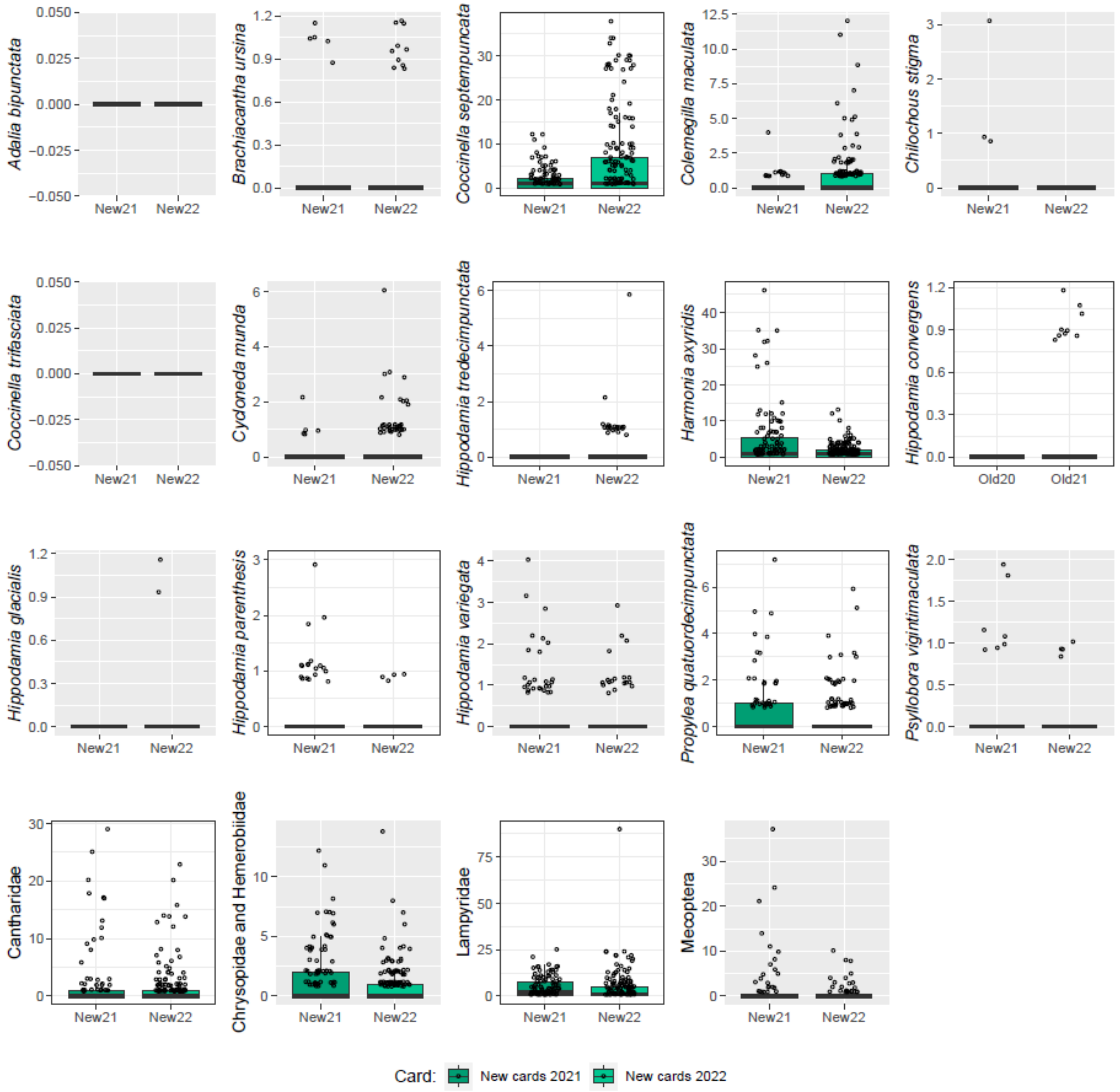


Figure S5: Captures of individual taxa for new cards in 2021 and 2022. Plots with a white background indicate significant difference in capture rates between year ($p < 0.05$). Plots with grey backgrounds indicate no statistical difference in capture rates were detected in this taxon.
